# Extended insight into the preponderance of LRRs - a docking & simulation study with members of the TLR1 subfamily

**DOI:** 10.1101/592626

**Authors:** Debayan Dey, Dipanjana Dhar, Sucharita Das, Aditi Maulik, Soumalee Basu

## Abstract

The widespread structural motif of Leucine-rich repeats (LRR) constitute the extracellular part of the Toll-like receptor (TLR) family preceded by an intracellular Toll/interleukin-1 receptor (TIR) domain at the C-terminus. The benefit of using LRRs in these pattern recognition receptors (PRR) that are responsible for early detection of pathogens to elicit inflammatory/innate immune response still remains elusive. Phylogenetic analyses (Maximum Likelihood and Bayesian Inference) of nine TLR (TLR 1-9) genes from 36 mammals reconfirmed the existence of two distinct clades, one (TLR1/2/6) for recognizing bacterial cell wall derivatives and another (TLR7/8/9) for various nucleic acids. TLR3, TLR4 and TLR5 showed independent line of evolution. The distinction of the TLR1 subfamily to form heterodimers within its members and the existence of the paralogs TLR1 and TLR6 therein, was appealing enough to carry out further studies with the extracellular recognition domain. Dimerizing and ligand binding residues from the crystal structures of TLR1 and TLR6 were interchanged to generate chimeric proteins. The dimer forming ability of these variants with their common partner, TLR2, were checked before running MD simulations. The chimeras were compared with wild type dimers to find no significant alterations in the overall structure. Finally, interchanged ligands were docked to the variants to ratify reversal of the binding function. Intriguingly, sequence change in substantial numbers, 16 in TLR1 and 18 in TLR6, preserves the native scaffold offered by LRRs. This exercise thus depicts how the LRR motif has been advantageous to be selected as an evolutionarily conserved motif for essential cellular processes.

## Introduction

Since its first revelation in 1985 as a repeating tetracosapeptide in the α_2_-glycoprotein of human serum [1], the evolutionarily conserved leucine-rich repeat (LRR) motifs have been found colossally in a number of proteins that encompass all the domains of life. The tandemly repeating motifs, that are 20-30 amino acid residues long, range from 2 to 62 in number and participate in a multitude of biologically significant processes [2-4]. Individual repeats are constituted of a leucine enriched highly conserved segment (HCS) consisting of a consensus sequence of 11 residues LxxLxLxxNxL or 12 residues LxxLxLxxCxxL and a variable segment (VS) [5]. Based on the variation of length and residue composition of the VS, LRRs can be categorized into 8 subclasses: ‘RI-like (Ribonuclease Inhibitor-like)’, ‘Cysteine-containing (CC)’, ‘bacterial’, ‘SDS22-like’, ‘Plant Specific (PS)’, ‘Typical’, ‘TpLRR (*Treponema pallidum*) LRR’ and ‘IRREKO’ [6]. The positioning of multiple repeats in tandem produces a solenoidal structure to provide a framework for protein-protein interaction. The concave surface of the solenoid is lined by parallel β-strands each of which correspond to HCS while the convex side is constituted of helical structures such that each helix correspond to the VS of individual LRRs [7, 8].

The importance of LRRs can be evaluated by the fact that mutations in human LRR-encoding genes have been linked with a number of diseases [9]. In addition, they participate in numerous crucial cellular processes like immunity, apoptosis, autophagy, cell polarization, neuronal development, nuclear mRNA transport, regulation of gene expression and ubiquitin-related processes [10, 11]. LRR proteins remain vastly expanded in immune repertoires of animals, mainly in invertebrates (e.g. sea urchin) and cephalochordates (e.g. *Amphioxus*), lacking adaptive immunity [12]. Both intracellular and extracellular LRRs exhibit functional diversity in the remarkably similar innate immune system of all organisms from plants to metazoans [13]. One of the most prominent examples of extracellular leucine-rich repeat constituting proteins are Toll-like receptors (TLRs), which prevail as a distinguishing representative of ‘Pattern Recognition Receptors’ (PRRs) for recognizing microbial molecular structures termed ‘Pathogen-associated Molecular Patterns’ (PAMPs).

Toll-like receptors, belonging to the type I transmembrane protein family, possess a tripartite domain organization with an N-terminal leucine-rich ectodomain (ECD), a central transmembrane region and a C-terminal Toll/interleukin-1 receptor (TIR) endodomain that mediates signalling pathways [14]. Although evolutionary studies reveal the presence of six TLR subfamilies (TLR1/2/6/10/14, TLR3, TLR4, TLR5, TLR7/8/9 and TLR11/12/13/21/22/23), not all TLR paralogs are expressed in different vertebrate species [15]. Despite formation of characteristic solenoidal structure in the extracellular portion and a conserved intracellular TIR domain, TLRs differ in ligand recognition through variable number and amino acid composition of LRRs. Quite a few crystal structures of TLR ectodomains complexed with their respective agonists namely TLR1-TLR2-triacylated lipopeptide, TLR2-TLR6-diacylated lipopeptide, TLR3-dsRNA, TLR4-MD2-lipopolysaccharide, TLR5-flagellin, TLR8-ssRNA, TLR9-CpG DNA and TLR13-ssRNA have been resolved [16-23]. Among the 10 functional TLRs in human, TLR2 exhibits distinctive ability of dimerizing with other members of TLR1 subfamily to elicit response against a myriad of membrane constituents derived from microbial pathogens [24]. Barring TLR2, TLR1/6/10 encoding genes are located consecutively on the same chromosome implying their emergence from an ancestral gene following successive gene duplication events. Although the heterodimers of TLR2/1 and TLR2/6 form very similar horseshoe shaped structures, the difference lies in the ligand binding as well as dimerizing residues of TLR1 and TLR6 that enable the expansion of their ligand repertoire [25].

Despite reports on the existence of preformed TLR2/1 and TLR2/6 heterodimers on the cell surface [26], recent findings confirmed the essentiality of ligands in stabilizing TLR2 heteromer interaction and activation. TLR2 in conjunction with TLR1 recognizes triacylated lipopeptide (Pam_3_CSK_4_) while TLR2/6 complex shows selective specificity towards diacylated lipopeptide (Pam_2_CSK_4_). Upon binding of TLR2/1 complex with Pam_3_CSK_4_, the 2 glycerol bound acyl chain fit into the hydrophobic groove of TLR2 and the third amide bound lipid chain into TLR1, thus bringing them into closer proximity for facilitating a stable interaction. TLR6, on the other hand, has a blocked lipid binding channel caused by the side chain of two phenylalanine residues (F343 and F365) and an approximately 80% increased exposed hydrophobic surface [27]. Both TLR1 and TLR6 are of similar length consisting of identical number of LRRs with the region LRR9-12 imparting the ability of differential lipopeptide recognition [28]. Hence, with so much in common between TLR1 and TLR6, it was highly intriguing to investigate the effect of interchanging functional residues between them keeping the native scaffold intact. This interchange if results in the reciprocation of the function would imply application of the LRR domain in protein engineering. In other words, desired change in function could be brought about through desired changes at specific positions of this domain. Variant proteins namely TLR1_6_ and TLR6_1_, were generated and docked to TLR2 for protein-protein interaction. Structural deformation at the monomer and dimer levels were analyzed throughout the molecular dynamics simulations carried out for the wild type and variant dimers. Finally, we docked the chimeras with reciprocal ligands to validate their functional ability.

## Results and Discussion

### Phylogeny of TLR1-9

Phylogenetic analyses (both Maximum Likelihood and Bayesian Inference) among the different organisms of the nine TLR members revealed individual clading pattern and highly identical tree topology with strong nodal support (Fig. 1 and Fig. 2) implying conservation of characteristic repeat numbers for each of the nine TLRs. Needless to say that the branching arrangements of the mammals in the tree followed the typical phylogenetic diversification of the TLR families, rather than leading to the intermixing of TLRs within closely related organisms. TLR1/2/6 formed a clade for recognizing bacterial cell wall derivatives whereas TLR7/8/9 that recognizes various nucleic acids clustered into another. TLR3, TLR4 and TLR5 showed independent line of evolution that justifies their differential ligand binding nature [29]. The nucleotide sequence of the Toll protein from the fruit fly (*Drosophila melanogaster*) was considered as outgroup for the study due to its structural and functional resemblance to mammalian Toll-like receptors [30].

**Fig. 1.**
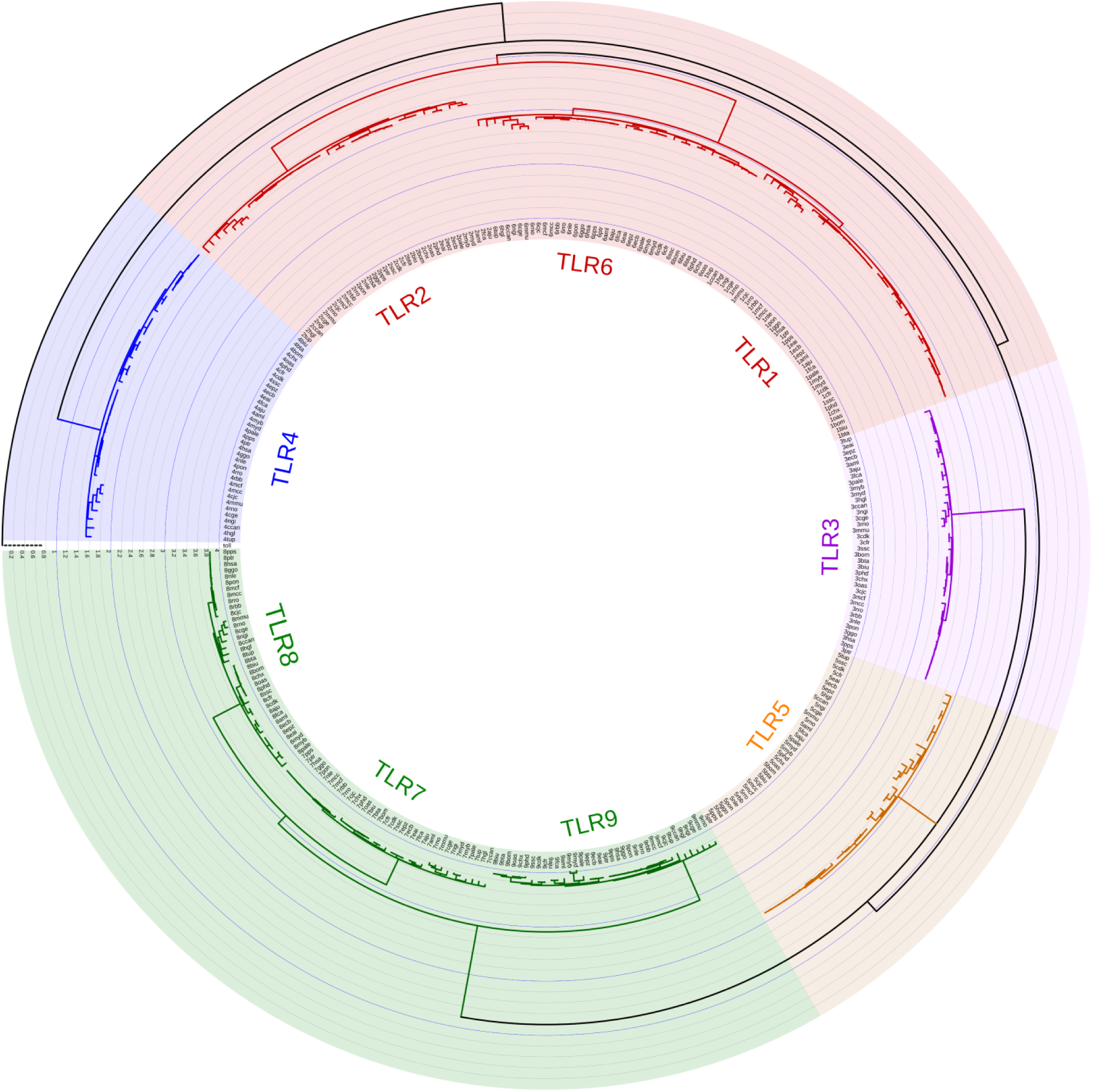
Maximum Likelihood tree of the nine TLRs from 36 mammals. The TLRs have been coloured according to the families. Only nodes with support values above 70% have been shown in the tree.

**Fig. 2.**
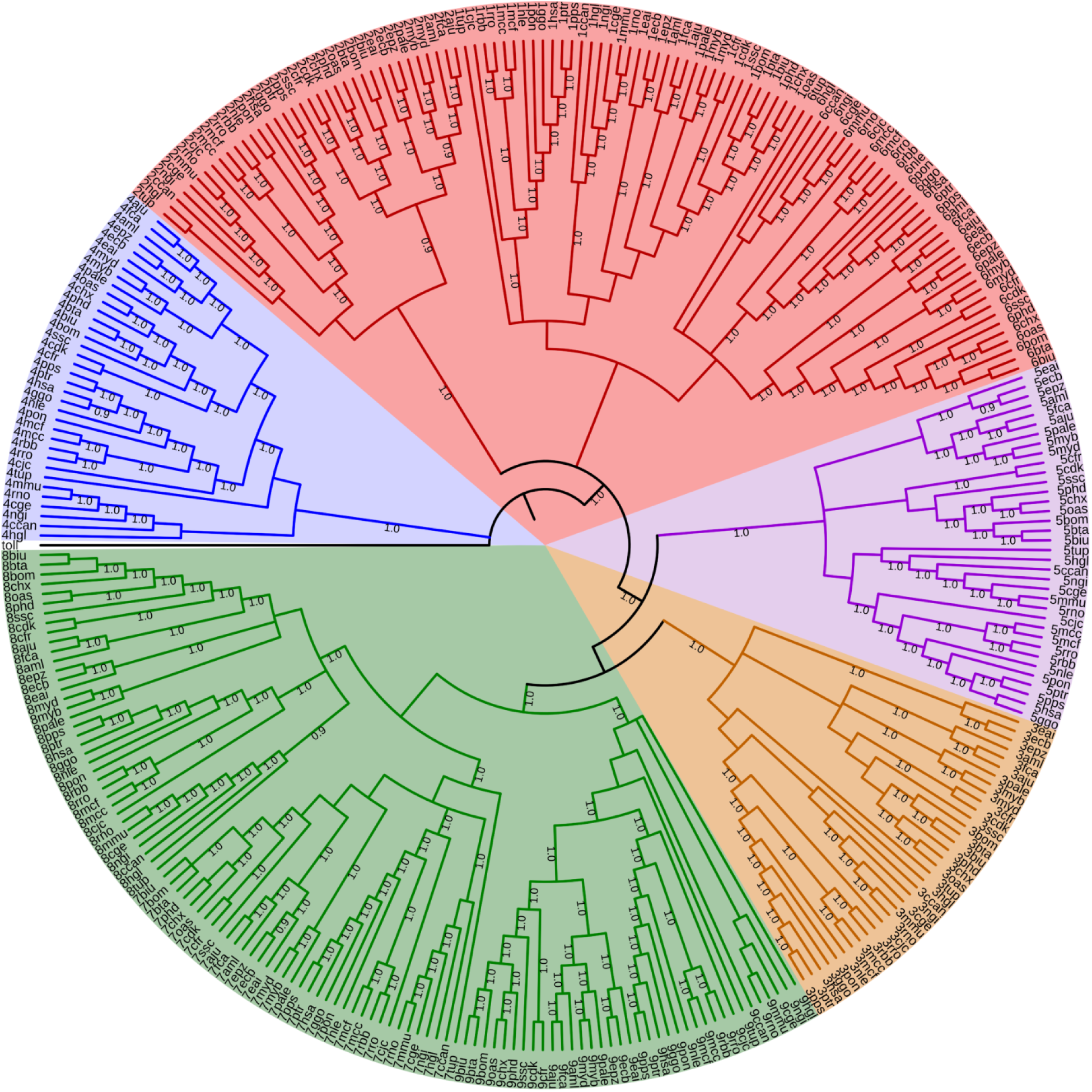
Bayesian Phylogeny of the nine TLRs from 36 organisms. The five TLR subfamilies considered are TLR1 (red), TLR3 (purple), TLR4 (blue), TLR5 (orange) and TLR7 (green). All nodes have more than 90% posterior probability.

### Structural Studies with TLRs

Among the different TLR subfamilies, the promiscuity of TLR2 lies in its ability to associate as heterodimers with other members of TLR1 subfamily (TLR1, TLR6, TLR10 and TLR14) for discerning the structurally broadest range of PAMPs [31]. Till today, TLR10 is characterized as an orphan receptor with no attributed ligands and remains as a disrupted gene in mice due to retroviral insertions [32]. Though TLR10 dimerizes with TLR2, it fails to initiate the classic TLR-associated signaling cascade whereas TLR14 remains exclusive to teleosts with unresolved function and ligand [15, 33]. Regardless of the fact that the heterodimerization of TLR2-TLR1 and TLR2-TLR6 enables the formation of distinct lipid-binding pockets to distinguish triacylated from diacylated lipopeptides, the subsequent immunological apparatus activate identical signaling pathways. Interestingly, each member of the dimers has 19 Leucine-rich repeats (LRRs) across the length of the extracellular segments that fold into a solenoidal curvature formed by uninterrupted array of tandem LRRs. Although the residues in the LRR9-12 regions of both TLR1 & TLR6 are responsible for agonist discrimination, an unusual high degree of conservation in the segment of 436-746 residues is observed implying a hetero-dynamic distribution of amino acids in these TLR scaffolds [34]. TLR1 & TLR6 being paralogs on the same chromosome, possess key amino acid residues that help in the identification of subtle differences between tri- and diacylated lipopeptides, upon dimerization with TLR2. The precise structural difference arises due to specific function imparting residues in the highly similar scaffold of these two heterodimers. Therefore, changes at the sequence level incorporated in the scaffold of the heterodimeric pair that would interchange their function with respect to ligand recognition might be the determining feature for binding specificity. To verify this, dimerizing and ligand binding residues of the TLRs (hTLR1 and mTLR6) were interchanged keeping the original scaffold of the proteins invariant.

### Structure based sequence alignment of TLR1 and TLR6

The crystal structure of human TLR2-TLR1 heterodimer bound to its natural ligand is available although the same for TLR2-TLR6 in ligand bound state from human is absent in PDB. On the same note, since the ligand bound complex of TLR2-TLR6 is available from mouse only, the following working strategy was adopted.

The structure based sequence alignment between hTLR1 and mTLR6 was implemented to determine the corresponding positions for incorporation of functionally interacting residues from TLR1 and TLR6. The dimerizing and ligand binding residues, for both hTLR1 and mTLR6, were marked in red and green, respectively while those possessing dual function (both dimerizing as well as ligand binding) were indicated in cyan. However, to identify the dimerizing and ligand binding residues of hTLR6 from mTLR6, the two sequences were aligned. Additionally, another pair of sequence was aligned to identify the functional residues of mTLR1 from hTLR1. In both the cases, the aligned residues at the corresponding positions were indicated using the same colour representation mentioned above.

In Fig. 3A, the residues of hTLR1 that were aligned with the marked residues (red, green and cyan) of mTLR6 were subjected to *in silico* mutation to the corresponding residue (Fig. 3B) at that position in the human homolog of TLR6 generating TLR1_6_ which is a variant TLR bearing the scaffold of TLR1 with dimerizing and ligand binding residues of TLR6. Likewise, in Fig. 4A, the residues of mTLR6 that were aligned with the marked residues of hTLR1 were targeted for *in silico* mutation to the corresponding aligned residues (Fig. 4B) in mouse homolog of TLR1 generating TLR6_1_. Additionally, two residues reported to be functionally important were substituted in the mTLR6_1_ (F343 and F365) with the corresponding amino acids of TLR1.

**Fig. 3.**
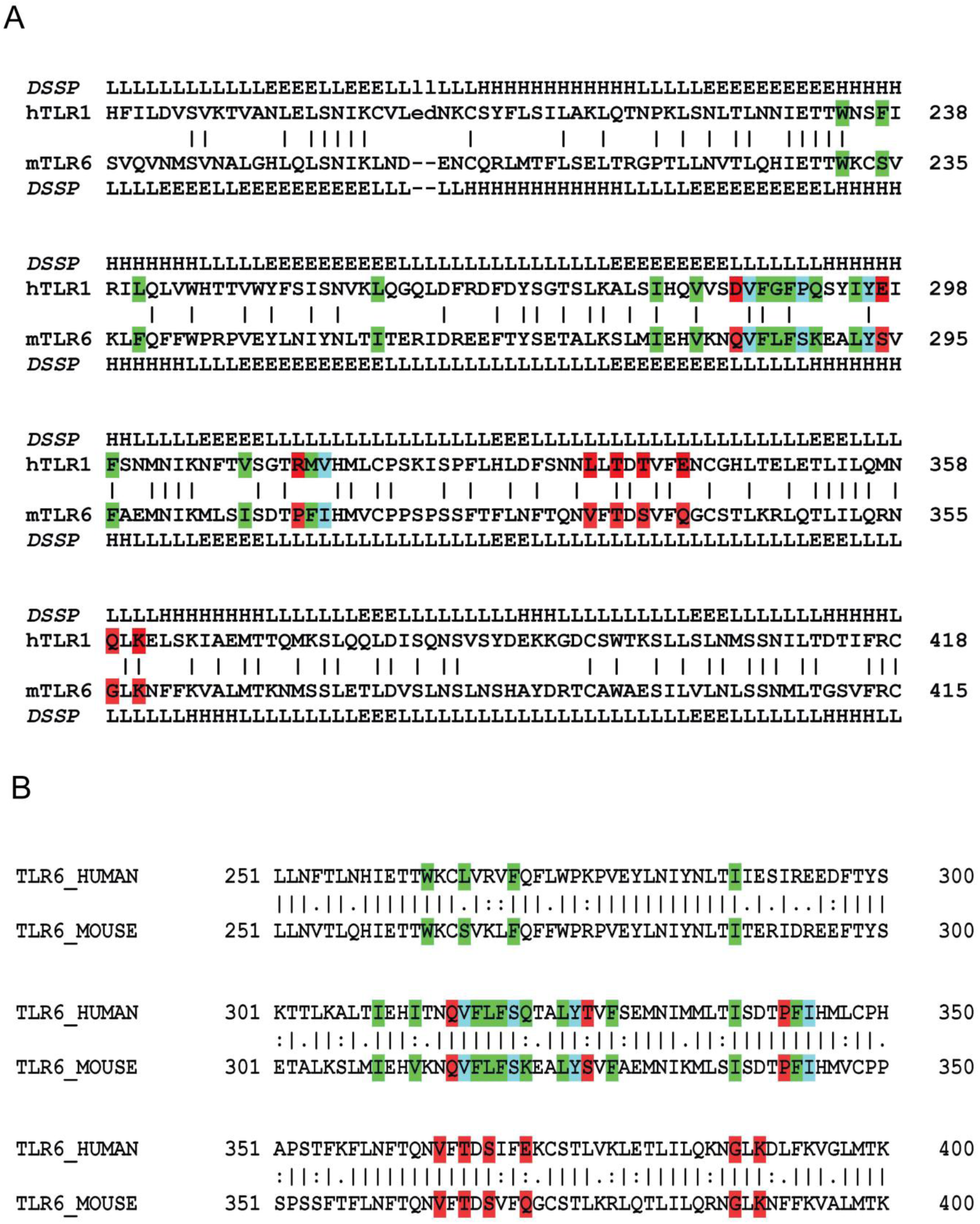
(A) Structural alignment of human TLR1 and mouse TLR6. Dimerizing and ligand binding residues of human TLR1 are marked along with the aligned residues in TLR6. (B) Pairwise sequence alignment of human TLR6 and mouse TLR6. The dimerizing and ligand binding residues were marked in red and green, respectively while those possessing dual function (both dimerizing as well as ligand binding) were indicated in cyan.

**Fig. 4.**
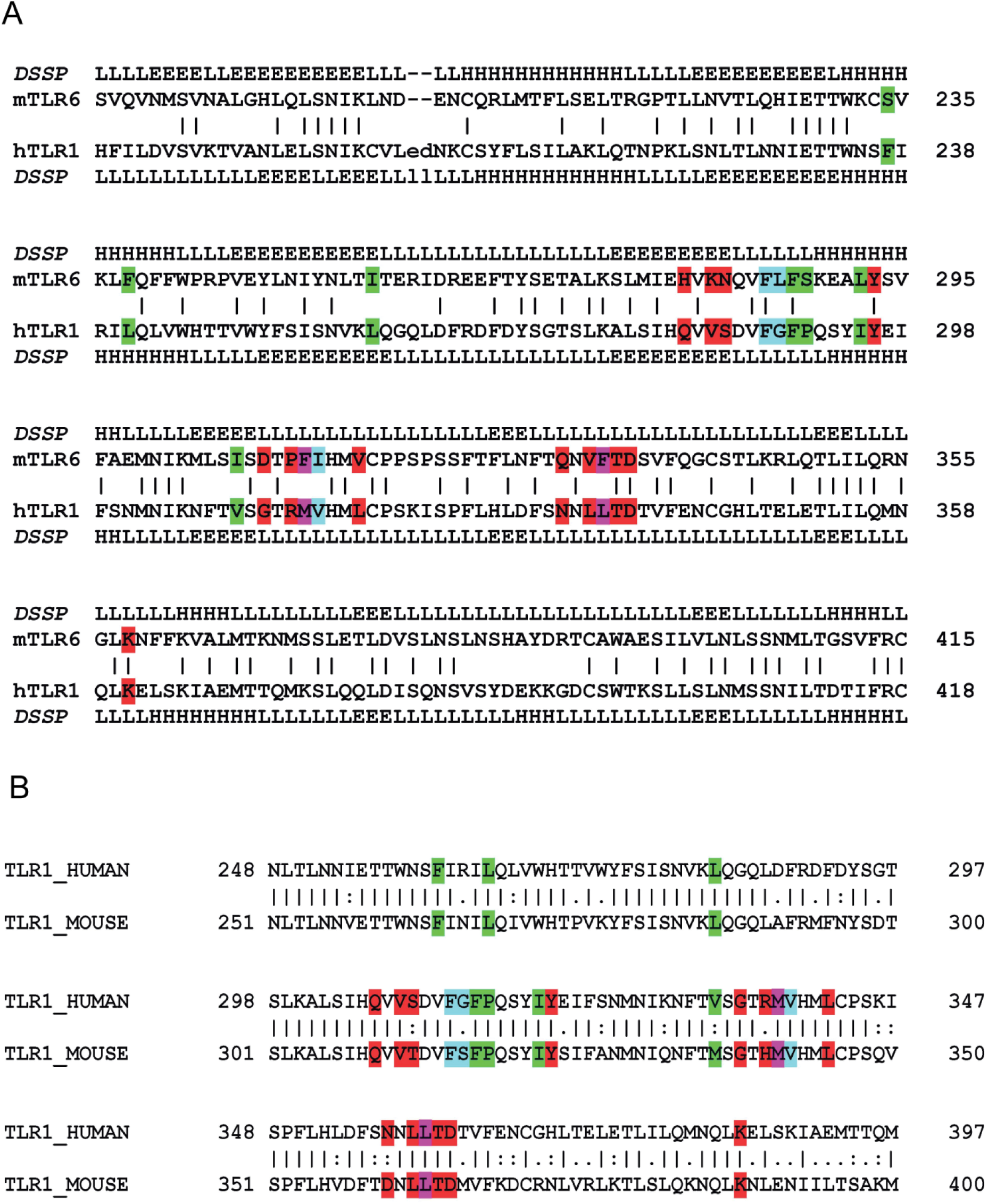
(A) Structural alignment of mouse TLR6 and human TLR1. Dimerizing and ligand binding residues of mouse TLR6 are marked along with the aligned residues in TLR1. Two functionally important residues are marked in magenta. (B) Pairwise sequence alignment of human TLR1 and mouse TLR1. The dimerizing and ligand binding residues were marked in red and green, respectively while those possessing dual function (both dimerizing as well as ligand binding) were indicated in cyan.

The obtained mutated monomers, hTLR1_6_ and mTLR6_1_, were subjected to energy minimization and individually docked with energy minimized structure of TLR2 (hTLR2 & mTLR2) giving rise to quite a few probable dimer structures of TLR1_6_-TLR2 from human and TLR6_1_-TLR2 from mouse. The potential dimer of TLR1_6_-TLR2 and TLR6_1_-TLR2 were selected from a group of structures sharing RMSD value of <4.5Å with the wild type dimer (Fig. 5). The selection of the mutated dimeric structures was validated through the change in Solvent Accessible Surface Area (SASA) of TLR1_6_ and TLR6_1_ using COCOMAPS (Table 1). The results showed significant changes in the residues of mutated monomers (hTLR1_6_ and mTLR6_1_) on dimerizing with their respective wild type protein (hTLR2 and mTLR2). In both the cases, five residues exhibited more than 70% change in the solvent accessible area upon dimerization.

**Table 1.**
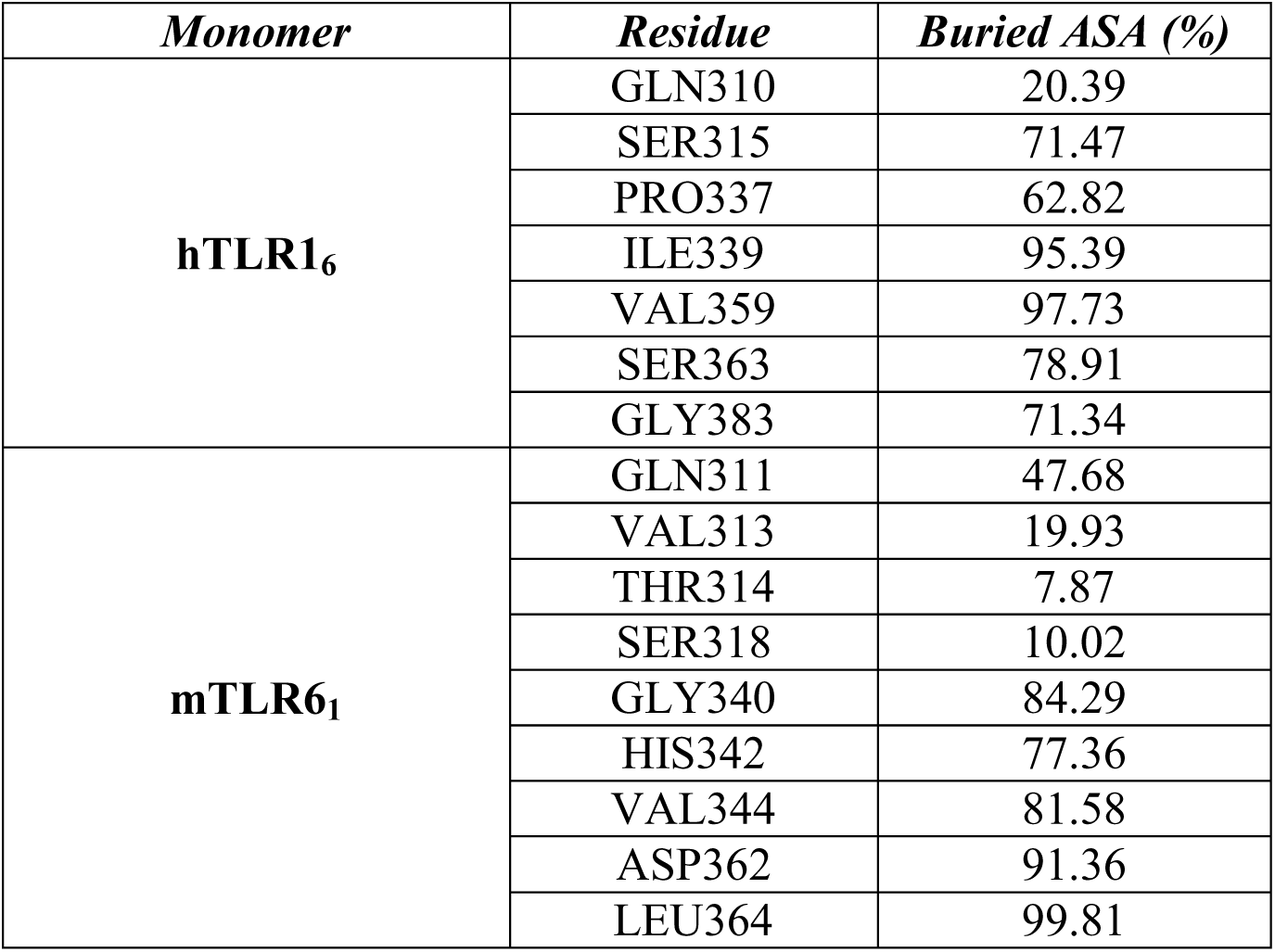
List of all the altered dimerizing residues of hTLR1_6_ and mTLR6_1_ showing change in Accessible Solvent Area (ASA).

| ***Monomer*** | ***Residue*** | ***Buried ASA (%)*** |
| --- | --- | --- |
| **hTLR1<sub>6</sub>** | GLN310 | 20.39 |
|  | SER315 | 71.47 |
|  | PRO337 | 62.82 |
|  | ILE339 | 95.39 |
|  | VAL359 | 97.73 |
|  | SER363 | 78.91 |
|  | GLY383 | 71.34 |
| **mTLR6<sub>1</sub>** | GLN311 | 47.68 |
|  | VAL313 | 19.93 |
|  | THR314 | 7.87 |
|  | SER318 | 10.02 |
|  | GLY340 | 84.29 |
|  | HIS342 | 77.36 |
|  | VAL344 | 81.58 |
|  | ASP362 | 91.36 |
|  | LEU364 | 99.81 |

**Fig. 5.**
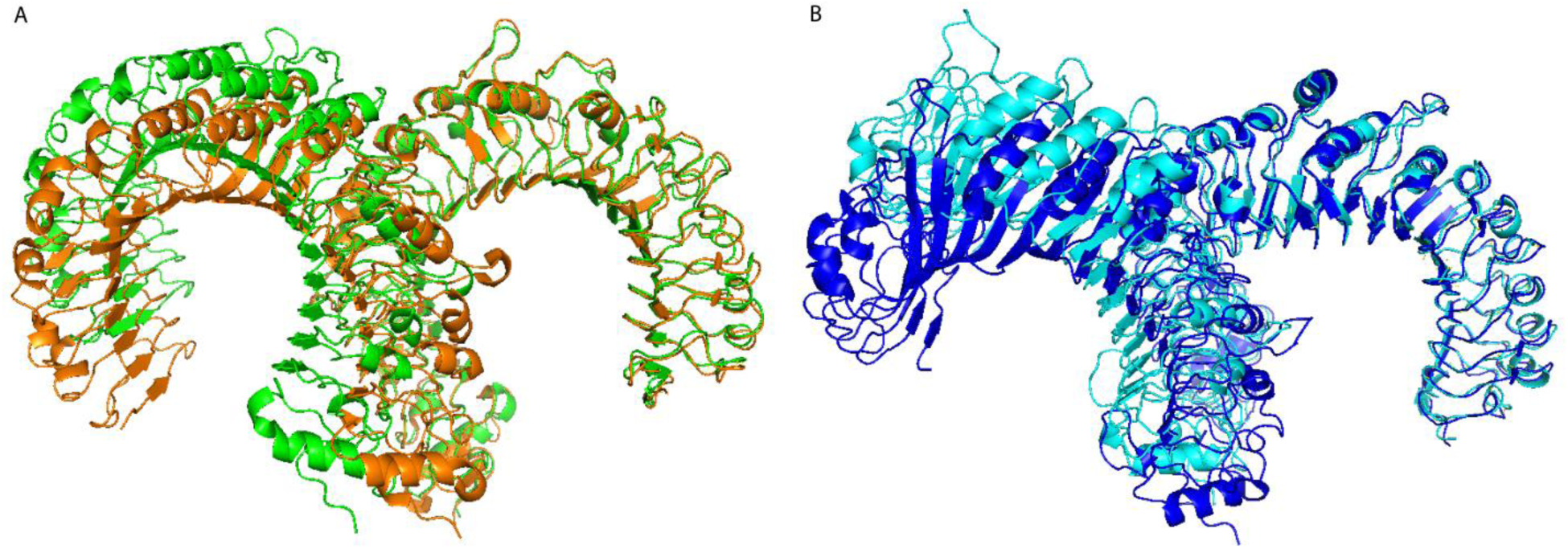
(A) Structures of hTLR1_6_-hTLR2 (orange) superimposed to hTLR1-hTLR2 (green). (B) Structures of mTLR6_1_-mTLR2 (cyan) superimposed to mTLR6-mTLR2 (blue).

### Molecular dynamics simulation analysis

Protein flexibility plays a significant role in TLRs for sensing invasion of pathogens and setting up an early innate immune response. Exchange of ligand binding and dimerizing residues between TLR1 and TLR6 has resulted in 16 and 18 changes respectively to each thus raising questions on the maintenance of the canonical structure of the two chimeras. We therefore conducted MD simulations of 60 ns on two wild type (hTLR1-hTLR2 & mTLR6-mTLR2) and two chimeric (hTLR1_6_-hTLR2 & mTLR6_1_-mTLR2) dimers for further analysis.

### Structural stability analysis

The time dependent changes of RMSD for the four heterodimers were assessed individually, considering the respective average structures as reference. The plots of RMSD reflected convergence of the simulations that are indicative of the overall protein stability (Fig. 6). The plots were depicted in two sets where the first graph corresponded to wild-type and variant hTLR1-hTLR2 and the other showing the same for mTLR6-mTLR2. It is evident that the time taken for both the chimeric dimers to reach convergence was more than their corresponding wild-type pairs. In Fig. 6A, both the wild-type (hTLR1-hTLR2) as well as the variant heteromers (hTLR1_6_-hTLR2) acquired stability from 20 ns. In Fig. 6B, the wild-type heterodimer of mTLR6-mTLR2 attained early stability owing to convergence from 25 ns whereas the mutated pair (mTLR6_1_-mTLR2) reached equilibrium much later.

**Fig. 6.**
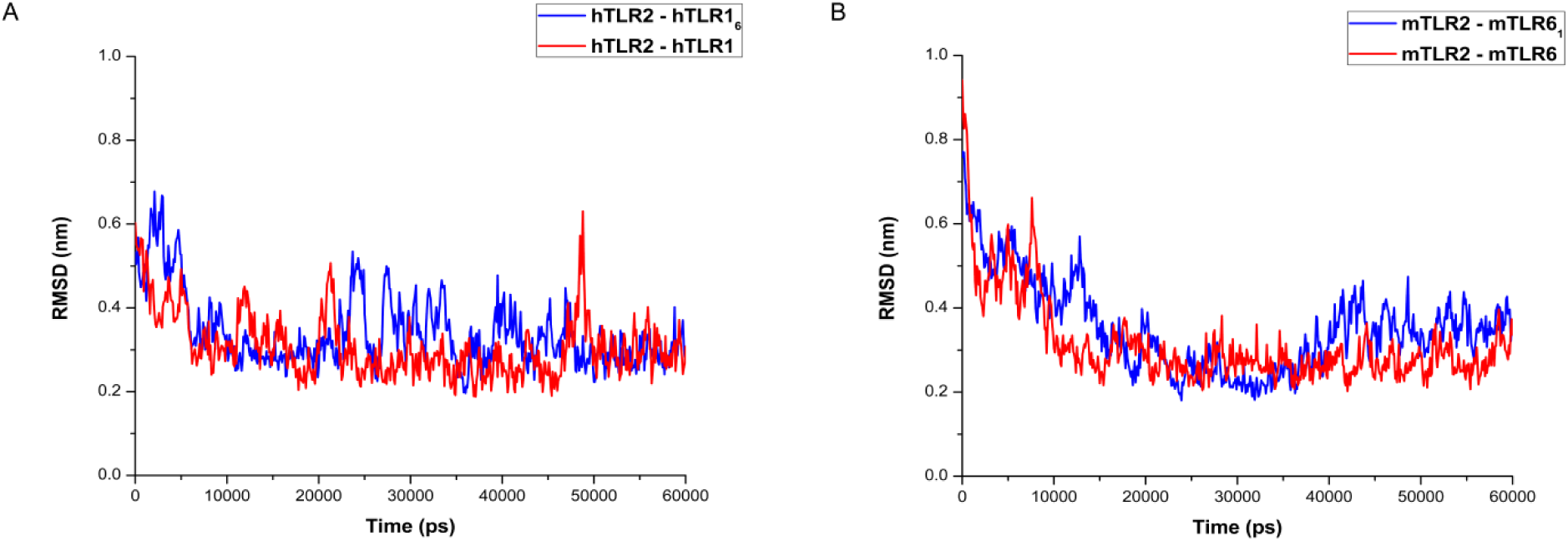
RMSD of (A) hTLR2-hTLR1_6_ & hTLR2-hTLR1 and (B) mTLR2-mTLR6_1_ & mTLR2-mTLR6.

### Comparison of protein flexibility between WT and mutated TLRs

The RMSF for individual residues for wild-type and mutated monomers were computed to infer the residue specific flexibility, taking into account the average structures as reference. In other words, four RMSF plots for hTLR1_6_ & hTLR1, mTLR6_1_ & mTLR6, hTLR2 (with hTLR1_6_ & hTLR1) and mTLR2 (with mTLR6_1_ & mTLR6) were generated (Fig. 7). It can be inferred that the C-terminal ends of both hTLR1_6_ and mTLR6_1_ showed fluctuations similar to their wild-type counterparts. Intriguingly, hTLR2 dimerized with hTLR1_6_, exhibited elevated flexibility at the central loop region and the C-terminal end whereas the former when partnered with mTLR6_1_ showed relatively less mobility at the central region. The minor difference in fluctuation patterns of hTLR2 and mTLR2 can be attributed to their subtle difference in the sequence and structural features [35]. However, in all the cases, the central portion was more restricted than the terminal regions owing to its functional importance. It is worth mentioning that in spite of possessing mutations in several significant residues on the otherwise unchanged TLR1 and TLR6 scaffolds, the RMSF values seemed to be in comparable concordance with their corresponding results of the wild-type monomers.

**Fig. 7.**
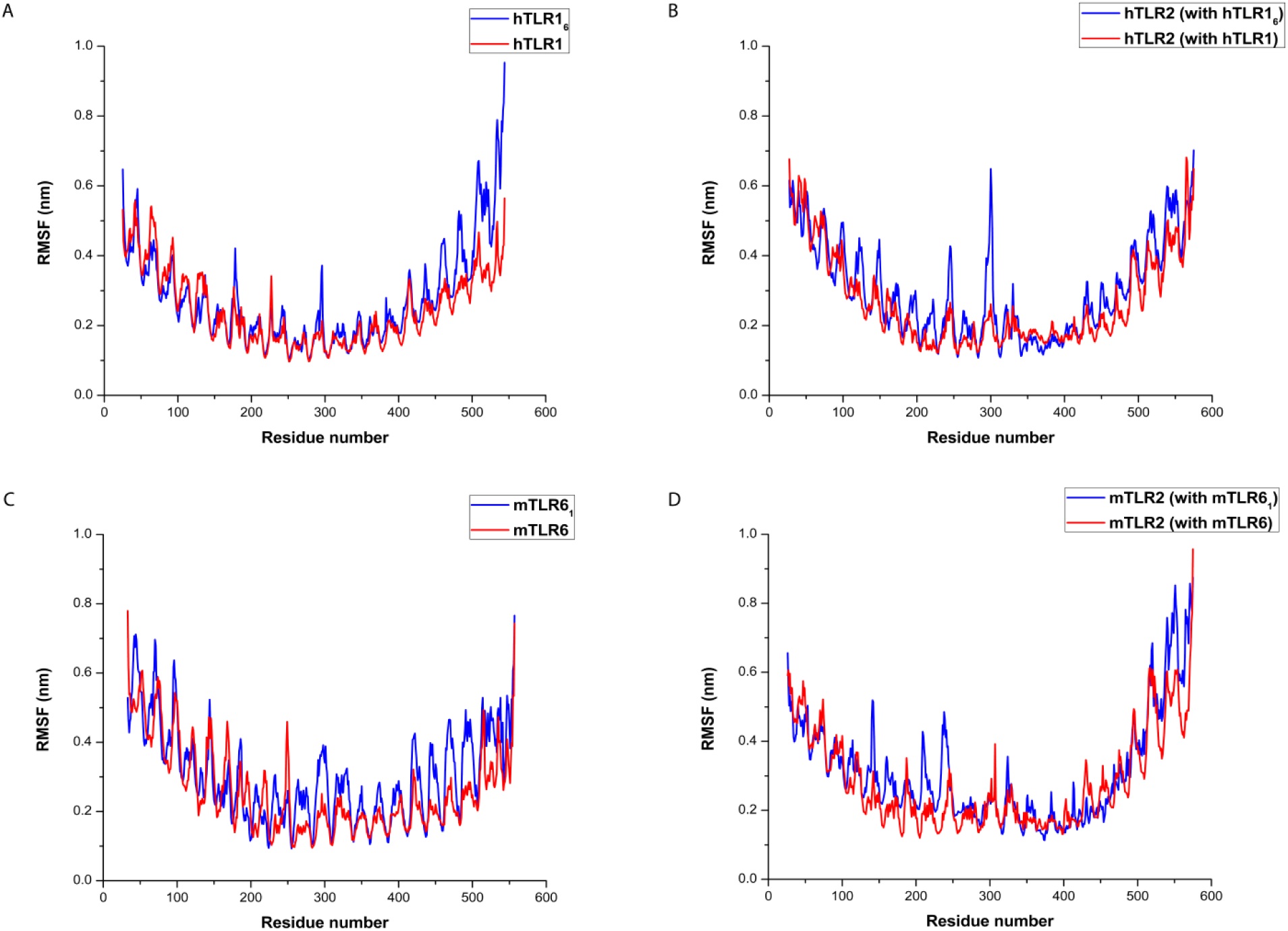
RMSF of (A) hTLR1 & hTLR16, (B) hTLR2 (with hTLR1 & hTLR1_6_), (C) mTLR6 & mTLR6_1_ and (D) mTLR2 (with mTLR6 & mTLR6_1_).

### Estimation of structural distortion

#### Variation in mouth width

The mouth width for both wild type and mutated TLR dimers was monitored individually for monomers, considering the center of mass (COM) of pair of terminal residues, to observe changes upon mutations in the protein that might lead to the opening or closing of the mouth. In general, we found that the mouth width of TLRs increased after simulation (Table 2). Comparison of the mouth width for the mTLR6 and mTLR6_1_ showed that the mouth width for the latter has been reduced (Fig. 8). It may be pointed out that mutated TLR6 (or TLR6_1_) has dimerizing and ligand binding residues of TLR1 on its own scaffold and therefore the range of variation of the mouth width matched with that of the hTLR1. Accordingly, mTLR2 bound to mTLR6_1_ showed a slight increase in mouth width when compared to the mTLR2 bound to mTLR6. This increase, as expected, was comparable to the hTLR2 bound to hTLR1. For the other dimer, although mutated TLR1 (or TLR1_6_) with dimerizing and ligand binding residues from TLR6, showed similar variation of mouth width as hTLR1, the range of variation was less than that of mTLR6. As expected, the mouth width variation of the hTLR2 bound to hTLR1_6_ is comparable to the variation of hTLR2 bound to mTLR6.

**Table 2.**
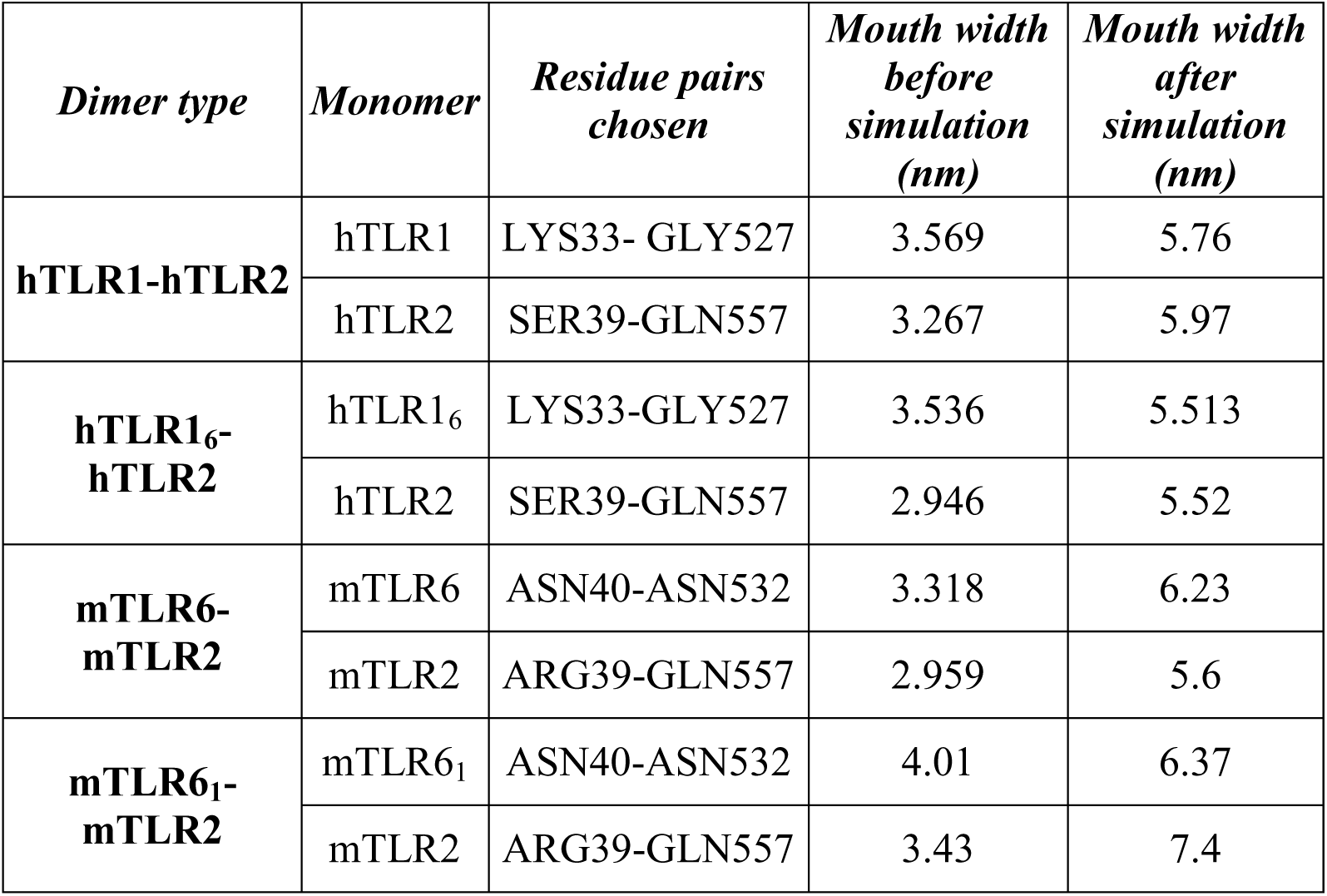
Comparison of the mouth width of the wild type and variant monomers before and after simulation.

**Fig. 8.**
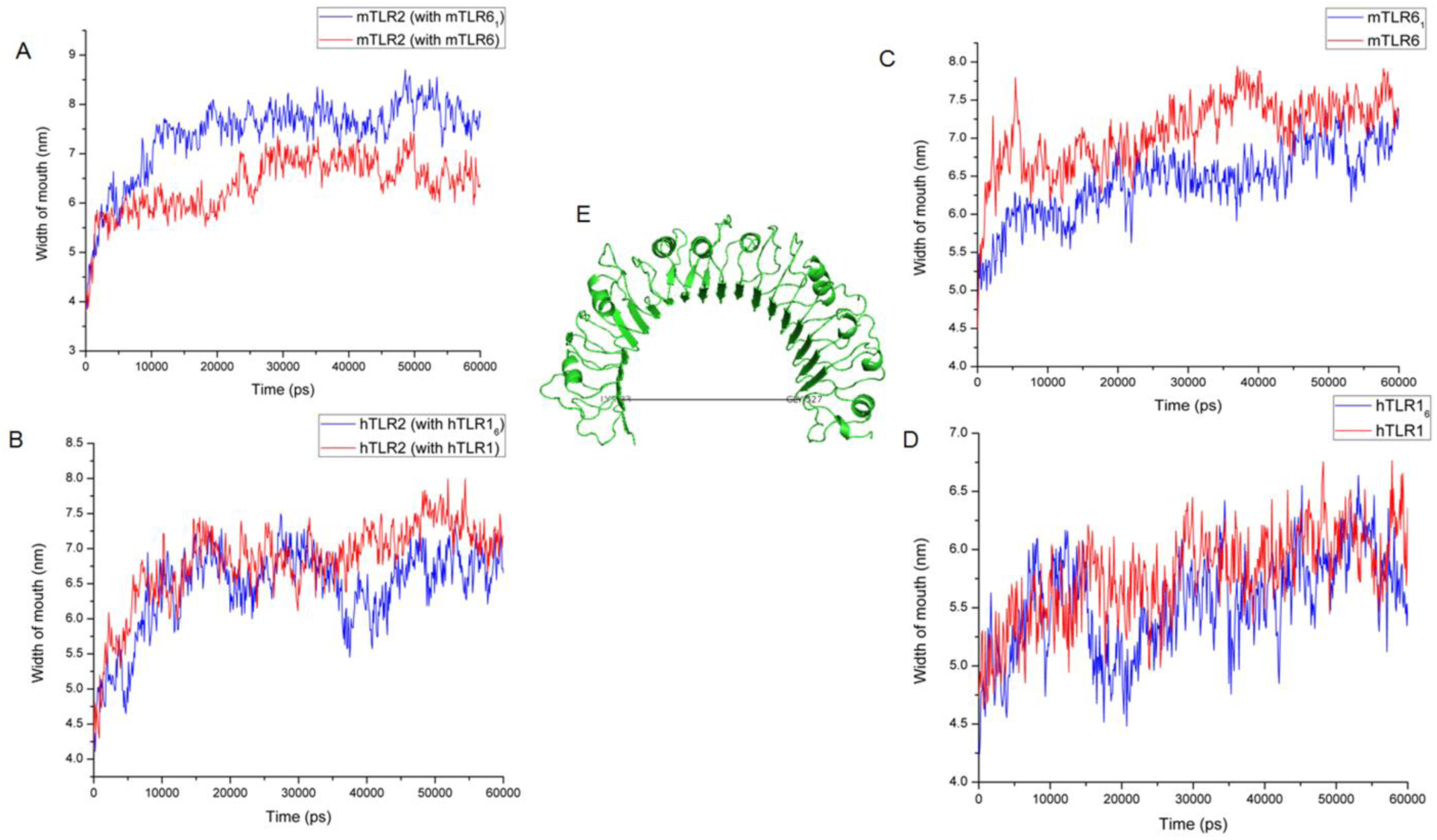
(A) Variation of mouth width of mTLR2 bound to mTLR6 and mTLR6_1_. (B) Variation of mouth width of hTLR2 bound to hTLR1 and hTLR1_6_. (C) Variation of mouth width of mTLR6 and mTLR6_1_. (D) Variation of mouth width of hTLR1 and hTLR1_6_. (E) Representation of mouth width.

#### Variation in the channel width

The lipid binding channel in hTLR1 is where one of the triacyl chains of Pam_3_CSK_4_ is housed. In hTLR1, the width of the mouth (TRP258-TYR320) and that at the central portion of the channel (TRP258-PHE323) was estimated and compared to the width of the mouth (TRP263-TYR325) and width of the central portion (TRP263-PHE328) in mTLR6 (Fig. 9). In the hTLR1_6_, the width of the mouth showed a slight increase than hTLR1. Additionally, the width of the central portion decreased in the mutated form. This might indicate that in hTLR1_6_ though the mouth of the channel had widened, the central region of the channel had narrowed. Therefore, as expected, hTLR1_6_ would no longer be able to accommodate the hTLR1 ligand. In case of mTLR6_1_ although the distance between the TRP263-TYR325 (mouth) showed an increase but that between TRP263-PHE328 (central region) remained almost unchanged when compared to the wild type. It may be noted that throughout the simulation, the width of the central region of mTLR6_1_ was higher than that of hTLR1 suggesting easy entry of the triacyl chain in the mTLR6_1_. Conclusively, estimation of the mouth and the channel width illustrated no significant structural deviation even upon incorporation of several amino acid changes.

**Fig. 9.**
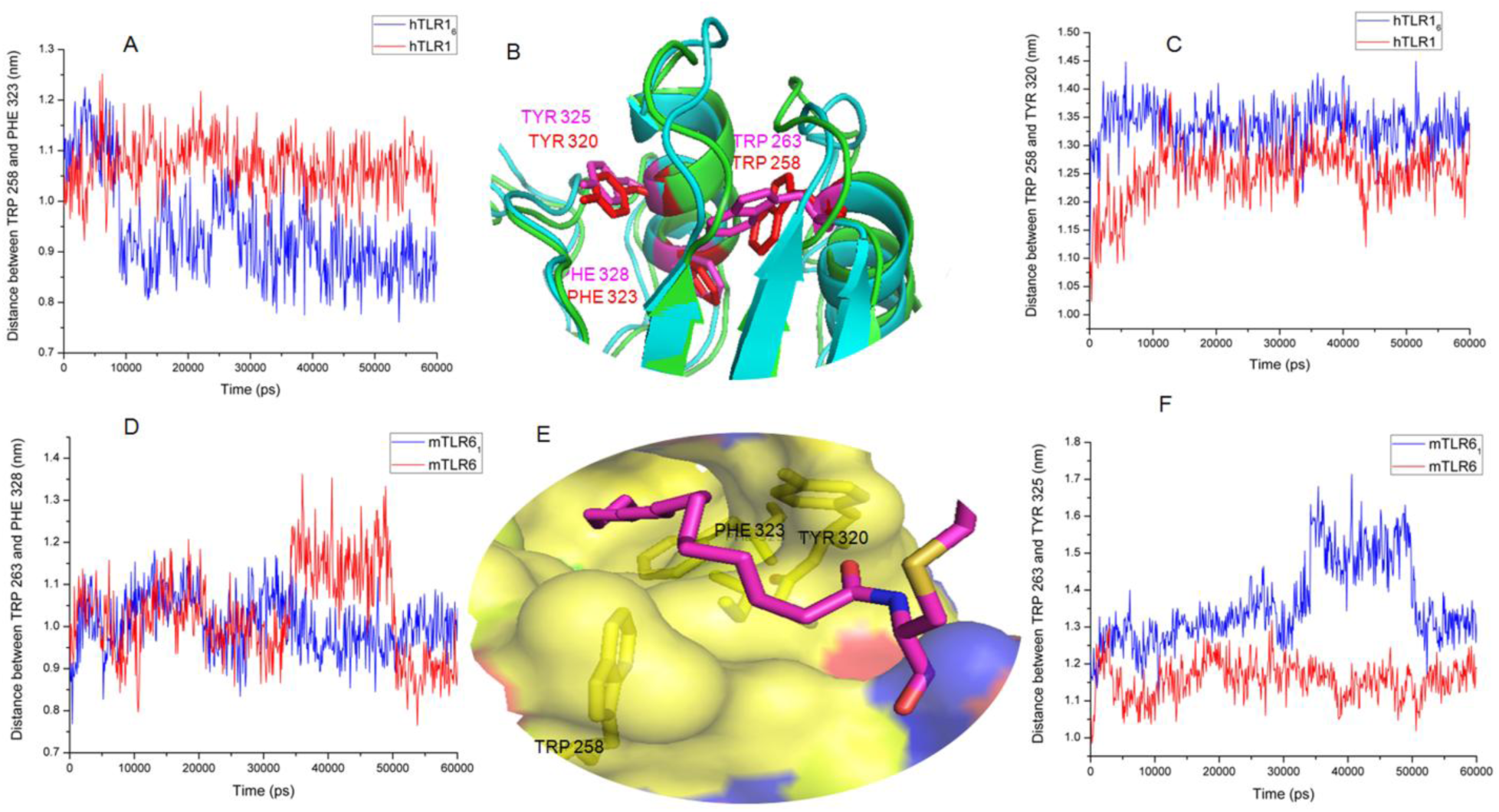
(A) Channel width of hTLR1 and hTLR1_6_ given by TRP-PHE distance. (B) Superimposed hTLR1 and mTLR6 channels. Channel lining residues are marked in red for hTLR1 and in magenta for mTLR6. (C) Channel width of hTLR1 and hTLR1_6_ given by TRP-TYR distance. (D) Channel width of mTLR6 and mTLR6_1_ given by TRP-PHE distance. (E) The hydrophobic channel of hTLR1 and the channel lining residues (F) Channel width of mTLR6 and mTLR6_1_ given by TRP-TYR distance.

### Ligand Docking Studies

The efficient utilization of TLR1 and TLR6 for differential dimerization with TLR2 enhances the recognition of a diverse ligand spectrum. Despite being paralogous in nature, subtle differences in the LRR9-12 regions of TLR1 and TLR6 allow the recognition of different ligands. At this juncture, we were interested in verifying whether the exchange of key residues amidst their native scaffolds resulted in reverse ligand recognition. The average simulated structures of hTLR1_6_-hTLR2 and mTLR6_1_-mTLR2 were docked to bacterial di- and triacylated lipopeptides, respectively, thus interchanging the natural ligands of the corresponding wild type scaffolds (Fig. 10). The variant dimers demonstrated the ability to bind the opposite ligands. We obtained favourable energy (−7.9 Kcal/mol for hTLR1_6_-hTLR2-diacylated lipopeptide and −7.8 Kcal/mol for mTLR6_1_-mTLR2-triacylated lipopeptide) and conformation for both the mutated pairs.

**Fig. 10.**
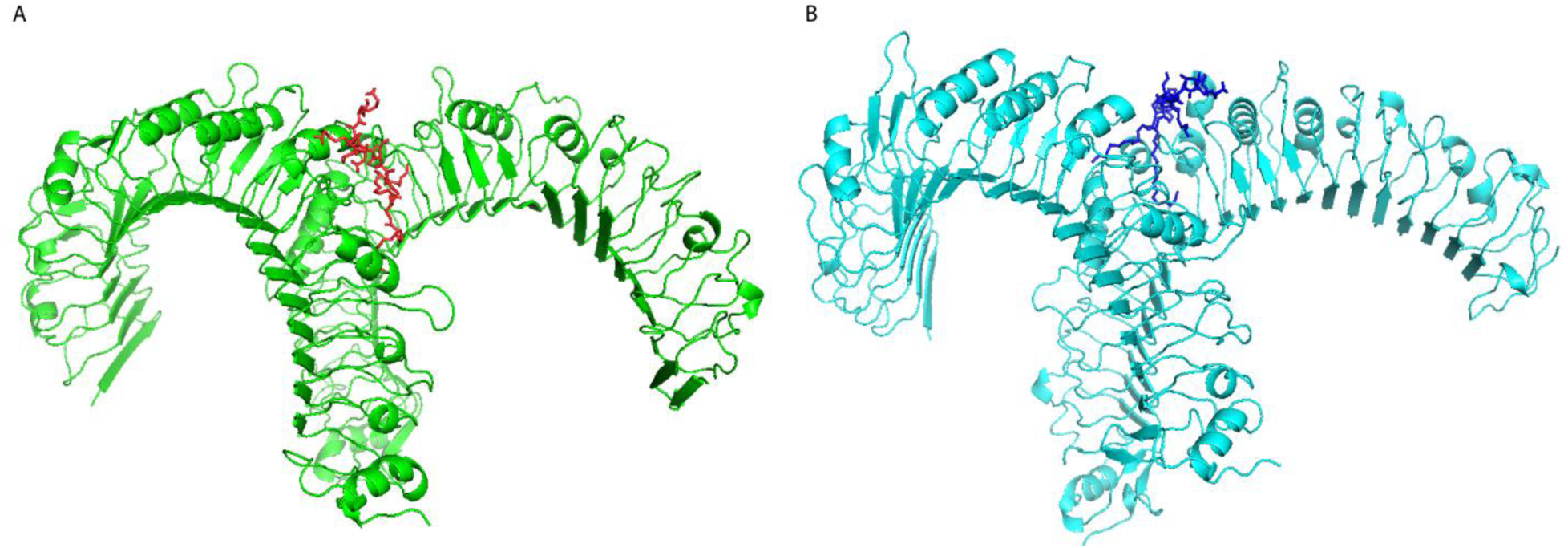
(A) Docked structure of hTLR1_6_-hTLR2 with diacylated lipopeptide. (B) Docked structure of mTLR6_1_-mTLR2 with triacylated lipopeptide.

## Materials and Methods

### Sequence retrieval

Nucleotide sequences of TLR1-9 for 36 organisms were retrieved from the KEGG GENES Database (Kyoto Encyclopedia of Genes and Genomes) (https://www.genome.jp/kegg/genes.html) [36] on July 14, 2018. The abbreviations for the organisms considered here, are listed as follows (Table 3):

**Table 3.** Three-letter codes and names of the organisms considered for phylogenetic studies.

|  | *Code* | *Scientific Name* | *Common Name* |
| --- | --- | --- | --- |
| 1 | hsa | *Homo sapiens* | Human |
| 2 | ptr | *Pan troglodytes* | Chimpanzee |
| 3 | pps | *Pan paniscus* | Bonobo |
| 4 | ggo | *Gorilla gorilla* | Western lowland gorilla |
| 5 | pon | *Pongo abelii* | Sumatran orangutan |
| 6 | nle | *Nomascus leucogenys* | Northern white-cheeked gibbon |
| 7 | mcc | *Macaca mulatta* | Rhesus monkey |
| 8 | mcf | *Macaca fascicularis* | Crab-eating macaque |
| 9 | rro | *Rhinopithecus roxellana* | Golden snub-nosed monkey |
| 10 | rbb | *Rhinopithecus bieti* | Black snub-nosed monkey |
| 11 | cjc | *Callithrix jacchus* | White-tufted-ear marmoset |
| 12 | mmu | *Mus musculus* | Mouse |
| 13 | rno | *Rattus norvegicus* | Rat |
| 14 | cge | *Cricetulus griseus* | Chinese hamster |
| 15 | ngi | *Nannospalax galili* | Upper Galilee mountains blind mole rat |
| 16 | hgl | *Heterocephalus glaber* | Naked mole rat |
| 17 | ccan | *Castor canadensis* | American beaver |
| 18 | tup | *Tupaia chinensis* | Chinese tree shrew |
| 19 | aml | *Ailuropoda melanoleuca* | Giant panda |
| 20 | fca | *Felis catus* | Domestic cat |
| 21 | aju | *Acinonyx jubatus* | Cheetah |
| 22 | bta | *Bos taurus* | Cow |
| 23 | bom | *Bos mutus* | Wild yak |
| 24 | biu | *Bos indicus* | Zebu Cattle |
| 25 | phd | *Pantholops hodgsonii* | Chiru |
| 26 | chx | *Capra hircus* | Goat |
| 27 | oas | *Ovis aries* | Sheep |
| 28 | ssc | *Sus scrofa* | Pig |
| 29 | cfr | *Camelus ferus* | Wild Bactrian camel |
| 30 | cdk | *Camelus dromedarius* | Arabian camel |
| 31 | ecb | *Equus caballus* | Horse |
| 32 | epz | *Equus przewalskii* | Przewalski's horse |
| 33 | eai | *Equus asinus* | Ass |
| 34 | myb | *Myotis brandtii* | Brandt's bat |
| 35 | myd | *Myotis davidii* | - |
| 36 | pale | *Pteropus alecto* | Black flying fox |

Only complete nucleotide sequences, corresponding to the full length of proteins, were considered. Each sequence possesses a characteristic KEGG accession ID [37]. It may be added that partial/precursor sequences were not taken into account for the study.

### Parameterization for Bayesian Analysis

The gene sequences obtained from the KEGG GENES Database were subjected to Multiple Sequence Alignment by ClustalW [38], with the default parameters. Thus, a total of 325 sequences (TLR1-9 from 36 organisms and an outgroup sequence ‘Toll’) were considered for the purpose. The output file of the alignment was saved in FASTA format for Bayesian inference.

The selection of appropriate nucleotide substitution models is an indispensable step for phylogenetic analysis of nucleotide sequence alignments. For accomplishment of the same, we used the software jModelTest2 (v2.1.10) [39]. jModelTest2 includes High Performance Computing (HPC) capabilities and additional features like new strategies for tree optimization, model-averaged phylogenetic trees (both topology and branch length), heuristic filtering and automatic logging of user activity [40]. According to jModelTest2, Transitional Model 3 + Proportion Invariant + Gamma (TIM3+I+G) was identified as the best-fit model for phylogenetic analyses using both corrected Akaike Information Criterion (cAIC) and Bayesian Information Criterion (BIC) calculations. Since this model could not be implemented in MrBayes 3.1.2, we replaced the former model with the closest parameterized model i.e. General Time Reversible + Proportion Invariant + Gamma (GTR+I+G) [41, 42].

### Phylogenetic Analysis

The aligned dataset was analyzed by Bayesian inference using MrBayes v3.2.6 [43] plugin in the Geneious R11.1 software (trial version), available at http://www.geneious.com [44]. A Markov chain Monte Carlo (MCMC) search with 2,000,000 generations was performed, logging results every 1000^th^ generation. The first 25% of the trees were discarded as burn-in. A consensus tree and the Bayesian Posterior Probabilities (BPP) were estimated based on the remaining trees. Bayesian inference was executed considering GTR+I+G nucleotide substitution model, which was calculated to be the most accurate evolutionary model by jModelTest2 (v2.1.10) according to both AIC and BIC.

Maximum likelihood phylogenetic analysis was also inferred using RAxML HPC2 Workflow on XSEDE (v8.2.10) through the CIPRES Science Gateway [45, 46]. The same nucleotide substitution model, *i.e.*, GTR+GAMMA+I was used for rate heterogeneity along with bootstrap analysis of 1000 generations. A majority rule consensus tree was derived from the 1000 bootstrapped trees. All the phylogenies generated by the aforementioned analyses were visualized using Figtree v1.4.3 (http://tree.bio.ed.ac.uk/software/figtree/) and iTOL v4.3 [47].

### Structural Studies involving TLRs

Crystal structures for dimer of human TLR2-TLR1 (PDB: 2Z7X) and mouse TLR2-TLR6 (PDB: 3A79) bound to their respective ligands, PAM_3_CSK_4_ and PAM_2_CSK_4_, in a definite stoichiometric ratio of 1:1 (heterodimeric protein complex: ligand) were obtained from the Protein Data Bank [16, 18]. The structural information of the ligand bound dimers was obtained from the PDB structures as follows:

1. **2Z7X**: Complex structure of TLR2 (Chain: A) - TLR1 (Chain: B) hetero-dimer from human bound with bacterial tri-acylated lipopeptide (Chain: C).
2. **3A79**: Complex structure of TLR2 (Chain: A) - TLR6 (Chain: B) hetero-dimer from mouse bound with bacterial di-acylated lipopeptide (Chain: C).

Pairwise structural alignment between TLR1 (PDB: 2Z7X, Chain: B) and TLR6 (PDB: 3A79, Chain: B) was performed using the Pairwise structure comparison tool in the DALI server (http://ekhidna2.biocenter.helsinki.fi/dali/). The DALI program computes optimal and suboptimal structural alignment, that is, a sequential set of one-to-one correspondences between C-alpha atoms [48]. The coordinates of the ligand binding and dimerizing residues, obtained from the crystal structures of both TLR2/1-PAM_3_CSK_4_ and TLR2/6-PAM_2_CSK_4_, were marked in separate structural alignments to identify the corresponding position in the aligned TLR for *in silico* mutagenesis. Owing to the unavailability of crystal structures for the ectodomain portion of human TLR6 and mouse TLR1, pairwise sequence alignments (https://www.ebi.ac.uk/Tools/psa/emboss_needle/) were carried out individually between human-mouse TLR6 and mouse-human TLR1 in order to maintain the structural correspondence at the species level [49]. Consequently, mutation of interacting residues of hTLR1 with hTLR6 and mTLR6 with mTLR1 resulted in the formation of two separate mutated monomers, hTLR1_6_ and mTLR6_1_ respectively.

Residues in one TLR scaffold were substituted by the dimerizing and ligand binding residues of another TLR using PyMOL [50], a Python-enhanced molecular graphics tool that specializes in 3D visualization of proteins, small molecules, density, surfaces, and trajectories. The *in silico* mutation was implemented using the Mutagenesis wizard by selecting the most probable rotamers for each amino acid substitution. Energy minimization was performed for each of the mutated (hTLR1_6_ and mTLR6_1_) and the wild-type (hTLR2 and mTLR2) monomeric TLR structures in vacuum and then in water using the GROMACS simulation package [51, 52]. The energy minimized structures were used for subsequent studies.

Protein-protein docking was performed using the ClusPro server (https://cluspro.org/) to obtain TLR heterodimers (hTLR1_6_-hTLR2 and mTLR6_1_-mTLR2) from mutated (hTLR1_6_ or mTLR6_1_) and wild-type (hTLR2 or mTLR2) monomers for molecular dynamics simulations and ligand docking studies [53].Docking in ClusPro involves three main steps. First, PIPER, a rigid body docking program based on the Fast Fourier Transform (FFT) correlation approach is extended to use pairwise interaction potentials. Second, the 1000 lowest energy conformations are clustered using pairwise IRMSD (Interface Root Mean Square Deviation) as the distance measure, and the 30 most populated clusters are retained for refinement. Third, the stability of the clusters is ensured with the removal of steric clashes by energy minimization. Protein-protein interacting residues in the heterodimeric complexes (hTLR1_6_-hTLR2 and mTLR6_1_-mTLR2) were identified from change in Accessible Surface Area (ASA) upon complex formation using COCOMAPS (bioCOmplexes Contact MAPS) [54].

The MD simulation protocol was applied to both the wild-type (as obtained from PDB) and mutated TLR heterodimers without their respective ligands. All simulations were performed using the GROMACS simulation package version 4.5.6. GROMOS96 53a6 united atom force field was used to model the intramolecular protein interactions and the intermolecular interactions between the protein and solvent molecules [55]. Initially the energy of each system was minimized using 500 steps of the steepest descent algorithm followed by 20,000 steps of the Polak-Ribiere Conjugate Gradient method to remove the strain in the initial structures. The relaxed structures were immersed in a rhombic dodecahedron of Simple Point Charge water molecules with periodic boundary conditions in all directions [56]. A minimum distance between the protein and wall of the cell was set to 1 nm to prevent the interactions between them. The solvated systems were neutralized by the addition of sodium and chloride ions to each of the systems according to the table given below (Table 4). These were followed by the energy minimization in solvent with the same steps as in vacuum but with periodic boundary conditions.

**Table 4.** Number of ions and solvent atoms added to the individual heteromeric systems during energy minimization protocol.

| *System* | *Sodium ions (NA)* | *Chloride ions (CL)* | *Solvent atoms* |
| --- | --- | --- | --- |
| **hTLR1-hTLR2** | 211 | 207 | 69975 |
| **hTLR1<sub>6</sub>-hTLR2** | 220 | 217 | 72738 |
| **mTLR6-mTLR2** | 219 | 219 | 72803 |
| **mTLR6<sub>1</sub>-mTLR2** | 206 | 205 | 69428 |

MD simulation studies consist of equilibration and production phases. In the first stage of equilibration, the solutes (protein, counter ions) were fixed, and the solvent (water molecules) was equilibrated for 100 ps of MD at 200 K using an integral time step of 0.001 ps. During the equilibration phase (i.e., at the start of the simulation), velocity was assigned to the atoms using Maxwell distribution. The system was coupled to the heat bath and heated to 300 K in a short run (100 ps) with 0.001 ps time step in which the system was allowed to relax in the new condition. This was followed by a short simulation (100 ps) with pressure coupling at 1 atm. During this phase, the velocities were reassigned according to a Maxwell distribution at 300 K. Finally, the production phase of MD simulation was run keeping the temperature, pressure and number of molecules of the ensemble invariant. Production phase was continued upto 60 ns using 0.002 ps time step. Subsequent analyses that include RMSD and RMSF were performed using different programs of the GROMACS package over 60 ns trajectory of the production run. The average structures were obtained using the same trajectory. The same trajectory has been used to estimate the mouth width of the TLRs. The distance between the center of mass of residues (COM) towards the end of the solenoid has been considered as the mouth width, calculated using gromacs. In case of mTLR2 (both with mTLR6 and mTLR6_1_) the residue pair chosen was ARG39-GLN557. Residue pair chosen for mTLR6 and mTLR6_1_ was ASN40-ASN532. For hTLR2 (both with hTLR1 and hTLR1_6_) the residue pair was SER 39-GLN 557. The residue pair chosen for hTLR1 and hTLR1_6_ was LYS33-GLY527. Additionally, residues lining the channel where one of the hydrophobic tails of the triacylated lipopeptide enters in TLR1 were identified. Thereafter, the corresponding TLR6 residues were identified by superposition in PyMOL. The distance between COM of two oppositely facing and channel lining residue pairs: (TRP258 - TYR320) and (TRP258 - PHE323) in TLR1 were used to obtain the channel width. The corresponding pairs for TLR6 include TRP263 - TYR325 and TRP263 - PHE328. The RMSD, RMSF, mouth width and channel width were visualized in the form of graphical representations with Origin8. The same residues were used to estimate the channel width for the mutated TLRs (hTLR1_6_ and mTLR6_1_).

The structure of the ligands were retrieved from PDB files (2Z7X and 3A79) for docking studies with the wild-type and mutated heterodimers of TLRs. Prior to this, the ligands were subjected to geometry optimization under the semi-empirical method in HyperChem™ 8.0.8 Molecular Modeling Software (Hypercube Inc., Gainesville, FL, USA). Both the Steepest Descent followed by Polak-Ribiere Conjugate Gradient algorithm was performed for energy optimization of PAM_3_CSK_4_ and PAM_2_CSK_4_ until convergence was reached. Open Babel was used for the interconversion of structures with different file formats [57]. Protein-ligand docking studies considering the mutated heterodimers with ligands were carried out using AutoDock Vina v1.1.2 [58]. The average structures obtained from MD simulation studies were subjected to docking with their respective ligands for determining their binding affinities. The pre-docking parameters were set using AutoDock Tools v4 [59] with the addition of hydrogen atoms and charges to the protein moieties. Protein and ligand files in the PDBQT format were used as input for molecular docking. The interactive visualization and images of all the protein structures were generated using PyMOL.

## Conclusion

Leucine-rich repeats (LRR) constituting the extracellular portion of TLRs are an ideal building block of repeat proteins that have been predominantly used as a motif for protein-protein interaction. The characteristic solenoidal shape of the repeats in tandem (LRR domain) is constituted by conserved residues forming parallel β-sheets along the concave surface and helical elements along the convex surface. Interestingly, such a protein domain is although likely to be activated by PAMPs for antimicrobial immune responses, it may be both extracellular and intracellular. This evolutionarily conserved domain is known to exhibit promiscuous cellular functions albeit innate immune response remains one of the dominant functions. Proteins containing LRRs have been extensively studied through several decades, yet, not much of our understanding unravels the advantage of its abundant usage.

Herein, the extracellular LRR domains of human TLR1, mouse TLR6 and their respective common dimerizing partner, TLR2, have been chosen for the study. It needs to be mentioned that all the three proteins are of comparable length and bear identical number of repeats. After dimer formation, whereas hTLR2-hTLR1 binds triacylated lipopeptide, mTLR2-mTLR6 binds diacylated lipopeptide. The binding specificity exhibited by the dimers with one different monomeric partner intrigued us to interchange the ligand-binding and the dimerizing residues between TLR1 and TLR6 and observe if reversal of function occurs. In other words, we wanted to observe whether reciprocating the ligand binding and dimerizing residues between TLR1 and TLR6 also reciprocates the binding of the ligands; triacylated lipopeptide and diacylated lipopeptide. The resulting chimeras had the scaffold of one TLR within which ligand binding and dimerizing residues of another from the same species (using pairwise alignment) were lodged (TLR1_6_, TLR6_1_). Combining the results of sequence alignment and docking with that of MD simulations we could conclude that the scaffold remaining that of TLR1, the presence of dimerizing and ligand binding residues of TLR6 (TLR1_6_) could make this variant TLR dimerize with TLR2 (in a similar manner as TLR6) and consequently bind diacylated lipopeptide, the cognate ligand of the TLR2-TLR6 dimer. Likewise the same holds true for the other TLR variant, TLR6_1_. Thus the structural stability imparted by the tandem LRRs could be realized through this exercise wherein the high number of changes made in the TLRs (16 in TLR1 and 18 in TLR6) destabilized neither the monomers nor the dimers throughout the simulations. Needless to say that favourable recognition/binding ability was observed through interchange of ligands. It is worth mentioning at this point that 9 out of the 16 changes in TLR1_6_ fall in the HCS of the LRRs and an even higher number of 14 among 18 fall in the same for TLR6_1_. To summarize, the elegance of this domain lies in the fact that it provides a scaffold for molecular interaction that is structurally robust to changes leaving ample scope for evolving with the interacting partner and thus effecting in unaltered recognition/ interaction for some cases or widening the repertoire of interacting molecules for others. The almost ubiquitous presence of the motif across wide range of cellular processes could thus be appreciated. Furthermore, identification of many single nucleotide polymorphisms in various TLR genes has been associated with particular diseases. Recently, several therapeutic agents targeting TLRs are under clinical and preclinical trials. This effort would add to insightful knowledge in this area apart from imparting fundamental understanding about the implications of the fascinating LRR motifs.

## Abbreviations

hTLR1: human Toll-like receptor1
hTLR2: human Toll-like receptor2
mTLR6: mouse Toll-like receptor6
mTLR2: mouse Toll-like receptor2
PAM_3_CSK_4_: synthetic triacylated lipopeptide
PAM_2_CSK_4_: synthetic diacylated lipopeptide
TLR1_6_: chimeric protein bearing the scaffold of TLR1 with dimerizing and ligand binding residues of TLR6
TLR6_1_: chimeric protein bearing the scaffold of TLR6 with dimerizing and ligand binding residues of TLR1.

## Acknowledgements

The authors are indebted to the Centre for High-Performance Computing for Modern Biology in the University of Calcutta. The authors also extend their sincere gratitude to the Bioinformatics Resources and Applications Facility (BRAF) of C-DAC, India, for providing supercomputing facility to execute long molecular dynamics simulations. We gratefully acknowledge Dr. Subinit Roy of Saha Institute of Nuclear Physics, Kolkata, for allowing us to use ORIGIN 8.0.

## References

1 Takahashi N, Takahashi Y & Putnam G (1985) Periodicity of leucine and tandem repetition of a 24-amino acid segment in the primary structure of leucine-rich alpha 2-glycoprotein of human serum. Proc. Natl. Acad. Sci. USA 82, 1906–1910.

2 Kobe B. & Deisenhofer J (1994) The leucine-rich repeat: a versatile binding motif. Trends in Biochemical Sciences 19, 415–421.

3 Buchanan SGSC & Gay NJ (1996) Structural and functional diversity in the leucine-rich repeat family of proteins. Progress in Biophysics and Molecular Biology 65, 1–44.

4 Matsushima N, Miyashita H, Mikami T & Kuroki Y (2010) A nested leucine rich repeat (LRR) domain: The precursor of LRRs is a ten or eleven residue motif. BMC Microbiology 10:235.

5 Kajava AV, Vassart G & Wodak SJ (1995) Modeling of the three-dimensional structure of proteins with the typical leucine-rich repeats. Structure 3, 867–877.

6 Kobe B & Kajava AV (2001) The leucine-rich repeat as a protein recognition motif. Curr. Opin. Struct. Biol 11, 725–732.

7 Kajava AV (1998) Stuctural diversity of leucine-rich repeat proteins. J. Mol. Biol. 277, 519–527.

8 Bella J, Hindle KL, McEwan PA & Lovell SC (2008) The leucine-rich repeat structure. Cell. and Mol. Life Sci 65, 2307–2333.

9 Matsushima N, Tachi N, Kuroki Y, Enkhbayar P, Osaki M, Kamiya M & Kretsinger RH (2005) Structural analysis of leucine-rich repeat variants in proteins associated with human diseases. Cell. Mol. Life Sci. 62, 2771–2791.

10 de Wit J, Hong W, Luo L & Ghosh A (2011) Role of leucine-rich repeat proteins in the development and function of neural circuits. Annu. Rev. Cell Dev. Biol. 27, 697–729.

11 Ng A & Xavier RJ (2011) Leucine-rich repeat (LRR) proteins: integrators of pattern recognition and signalling in immunity. Autophagy 7, 1082–1084.

12 Huang S, Yuan S, Guo L, Yu Y, Li J, Wu T, Liu T, Yang M, Wu K, Liu H, Ge J, Yu Y, Huang H, Dong M, Yu C, Chen S, & Xu A (2008) Genomic analysis of the immune gene repertoire of amphioxus reveals extraordinary innate complexity and diversity Genome Res. 18, 1112–1126.

13 Nürnberger T, Brunner F, Kemmerling B & Piater L (2004) Innate immunity in plants and animals: striking similarities and obvious differences. Immunological Reviews 198, 249–266.

14 Bell JK, Mullen GED, Leifer CA, Mazzoni A, Davies DR and Segal DM (2003) Leucine-rich repeats and pathogen recognition in Toll-like receptors. Trends in Immunology 24, 528–533.

15 Roach JC, Glusman G, Rowen L, Kaur A, Purcell MK, Smith KD, Hood LE & Aderem A (2005) The evolution of vertebrate Toll-like receptors. Proc. Natl. Acad. Sci. USA 102, 9577–9582.

16 Jin M, Kim S, Heo J, Lee M, Kim H, Paik S, Lee H & Lee J (2007) Crystal structure of the TLR1-TLR2 heterodimer induced by binding of a tri-acylated lipopeptide. Cell 130, 1071–1082.

17 Liu L, Botos I, Wang Y, Leonard JN, Shiloach J, Segal DM & Davies DR (2008) Structural basis of Toll-like receptor 3 signaling with double-stranded RNA. Science 320, 379–381.

18 Kang J, Nan X, Jin M, Youn S, Ryu Y, Mah S, Han S, Lee H, Paik S & Lee J (2009) Recognition of lipopeptide patterns by Toll-like receptor 2-Toll-like receptor 6 heterodimer. Immunity 31, 873–884.

19 Park BS, Song DH, Kim HM, Choi BS, Lee H & Lee JO (2009) The structural basis of lipopolysaccharide recognition by the TLR4–MD-2 complex. Nature 458, 1191–1195.

20 Yoon SI, Kurnasov O, Natarajan V, Hong M, Gudkov AV, Osterman AL & Wilson IA Structural basis of TLR5-flagellin recognition and signaling. Science 335, 859–864.

21 Tanji H, Ohto U, Shibata T, Taoka M, Yamauchi Y, Isobe T, Miyake K & Shimizu T (2015) Toll-like receptor 8 senses degradation products of single-stranded RNA. Nat. Struct. Mol. Biol. 22, 109–115.

22 Ohto U, Shibata T, Tanji H, Ishida H, Krayukhina E, Uchiyama S, Miyake K & Shimizu T (2015) Structural basis of CpG and inhibitory DNA recognition by Toll-like receptor 9. Nature 520, 702–705.

23 Song W, Wang J, Han Z, Zhang Y, Zhang H, Wang W, Chang J, Xia B, Fan S, Zhang D, Wang J Wang HW & Chai J (2015) Structural basis for specific recognition of single-stranded RNA by Toll-like receptor 13. Nature Structural & Molecular Biology 22, 782–787.

24 Ozinsky A, Underhill DM, Fontenot JD, Hajjar AM, Smith KD, Wilson CB, Schroeder L & Aderem A (2000) The repertoire for pattern recognition of pathogens by the innate immune system is defined by cooperation between toll-like receptors. Proc. Natl. Acad. Sci. USA 97, 13766–13771.

25 Botos I, Segal DM & Davies DR (2011) The structural biology of Toll-like receptors. Structure 19, 447–459.

26 Triantafilou M, Gamper FGJ, Haston RM, Mouratis MA, Morath S, Hartung T & Triantafilou K (2006) Membrane Sorting of Toll-like Receptor (TLR)-2/6 and TLR2/1 Heterodimers at the Cell Surface Determines Heterotypic Associations with CD36 and Intracellular Targeting. The Journal of biological chemistry 281, 31002–31011.

27 van Bergenhenegouwen J, Plantinga TS, Joosten LA, Netea MG, Folkerts G, Kraneveld AD, Garssen J & Vos AP (2013) TLR2 & Co: a critical analysis of the complex interactions between TLR2 and coreceptors. J Leukoc Biol. 94, 885–902.

28 Omueti KO, Beyer JM, Johnson CM, Lyle EA & Tapping IR (2005) Domain Exchange between Human Toll-like Receptors 1 and 6 Reveals a Region Required for Lipopeptide Discrimination. The Journal of biological chemistry 280, 36616–36625.

29 Dhar D, Dey D & Basu S (2019) Insights into the evolution of extracellular leucine-rich repeats in metazoans with special reference to Toll-like receptor 4. Journal of Biosciences 44.

30 Rock FL, Hardiman G, Timans JC, Kastelein RA & Bazan JF (1998) A family of human receptors structurally related to *Drosophila* Toll. Proceedings of the National Academy of Sciences USA 95, 588–593.

31 Matsushima N, Tanaka T, Enkhbayar P, Mikami T, Taga M, Yamada K & Kuroki Y (2007) Comparative sequence analysis of leucine-rich repeats (LRRs) within vertebrate toll-like receptors. BMC Genomics 8, 124.

32 Hasan U, Chaffois C, Gaillard C, Saulnier V, Merck E, Tancredi S, Guiet C, Brière F, Vlach J, Lebecque S, Trinchieri G & Bates EEM (2005) Human TLR10 Is a Functional Receptor, Expressed by B Cells and Plasmacytoid Dendritic Cells, Which Activates Gene Transcription through MyD88. J Immunol. 174, 2942–2950.

33 Guan Y, Ranoa DR, Jiang S, Mutha SK, Li X, Baudry J & Tapping RI (2010) Human TLRs 10 and 1 share common mechanisms of innate immune sensing but not signaling. J Immunol. 184, 5094–5103.

34 Kruithof EKO, Satta N, Liu JW, Dunoyer-Geindre S & Fish RJ (2007) Gene conversion limits divergence of mammalian TLR1 and TLR6. BMC Evolutionary Biology 7,148.

35 Durai P, Rajiv Gandhi Govindaraj RG & Choi S (2013) Structure and dynamic behavior of Toll-like receptor 2 subfamily triggered by malarial glycosylphosphatidylinositols of *Plasmodium falciparum*. The FEBS Journal 280, 6196–6212.

36 Kanehisa, Furumichi M, Tanabe M, Sato Y & Morishima K (2017) KEGG: new perspectives on genomes, pathways, diseases and drugs. Nucleic Acids Res 45, D353–D361.

37 O’Leary NA, Wright MW, Brister JR, Ciufo S, Haddad D, McVeigh R, Rajput B, Robbertse B, Smith-White B, Ako-Adjei D, Astashyn A, Badretdin A, Bao Y, Blinkova O, Brover V, Chetvernin V, Choi J, Cox E, Ermolaeva O, Farrell CM, Goldfarb T, Gupta T, Haft D, Hatcher E, Hlavina W, Joardar VS, Kodali VK, Li W, Maglott D, Masterson P, McGarvey KM, Murphy MR, O’Neill K, Pujar S, Rangwala SH, Rausch D, Riddick LD, Schoch C, Shkeda A, Storz SS, Sun H, Thibaud-Nissen F, Tolstoy I, Tully RE, Vatsan AR, Wallin C, Webb D, Wu W, Landrum MJ, Kimchi A, Tatusova T, DiCuccio M, Kitts P, Murphy TD & Pruitt KD (2016) Reference sequence (RefSeq) database at NCBI: current status, taxonomic expansion, and functional annotation. Nucleic Acids Research 44, D733–745.

38 Sievers F, Wilm A, Dineen DG, Gibson TJ, Karplus K, Li W, Lopez R, McWilliam H, Remmert M, Söding J, Thompson JD & Higgins DG (2011) Fast, scalable generation of high-quality protein multiple sequence alignments using Clustal Omega. Molecular Systems Biology 7:539.

39 Posada D (2008) jModelTest: phylogenetic model averaging. Molecular Biology and Evolution 25, 1253–1256.

40 Darriba D, Taboada GL, Doallo R & Posada D (2012) jModelTest 2: more models, new heuristics and parallel computing. Nature Methods 9:772.

41 Lecocq T, Vereecken NJ, Michez D, Dellicour S, Lhomme P, Valterová I, Rasplus JY & Rasmont P (2013) Patterns of Genetic and Reproductive Traits Differentiation in Mainland vs. Corsican Populations of Bumblebees. PLoS One 8:e65642.

42 Arenas M (2015) Trends in substitution models of molecular evolution. Frontiers in Genetics 6:319.

43 Ronquist F, Teslenko M, van der Mark P, Ayres DL, Darling A, Höhna S, Larget B, Liu L, Suchard MA & Huelsenbeck JP (2012) MrBayes 3.2: Efficient Bayesian Phylogenetic Inference and Model Choice Across a Large Model Space. Systematic Biology 61, 539–542.

44 Kearse M, Moir R, Wilson A, Stones-Havas S, Cheung M, Sturrock S, Buxton S, Cooper A, Markowitz S, Duran C, Thierer T, Ashton B, Mentjies P & Drummond A (2012) Geneious Basic: an integrated and extendable desktop software platform for the organization and analysis of sequence data. Bioinformatics 28, 1647–1649.

45 Miller MA, Pfeiffer W & Schwartz T (2010) “Creating the CIPRES Science Gateway for inference of large phylogenetic trees”. Proceedings of the Gateway Computing Environments Workshop (GCE). New Orleans, LA 1–8.

46 Stamatakis A (2014) RAxML version 8: a tool for phylogenetic analysis and post-analysis of large phylogenies. Bioinformatics 30, 1312–1313.

47 Letunic I & Bork P (2016) Interactive tree of life (iTOL) v3: an online tool for the display and annotation of phylogenetic and other trees. Nucleic Acids Research 44, 242–245.

48 Holm L & Laakso LM (2016) Dali server update. Nucleic Acids Research 44, W351–W355.

49 Li W, Cowley A, Uludag M, Gur T, McWilliam H, Squizzato S, Park YM, Buso N & Lopez R (2015) The EMBL-EBI bioinformatics web and programmatic tools framework. 43, W580–584.

50 The PyMOL Molecular Graphics System, Version 1.7 Schrödinger, LLC.

51 Berendsen HJC, van der Spoel D & van Drunen R (1995) GROMACS: A message-passing parallel molecular dynamics implementation. Computer Physics Communications 91, 43–56.

52 van der Spoel D, Lindahl E, Hess B, van Buuren AR, Apol E, Meulenhoff PJ, Tieleman DP, Sijbers ALTM, Feenstra KA, van Drunen R & Berendsen HJC (2010) Gromacs User Manual version 4.5.6.

53 Kozakov D, Hall DR, Xia B, Porter KA, Padhorny D, Yueh C, Beglov D & Vajda S (2017) The ClusPro web server for protein-protein docking. Nature Protocols 12, 255–278.

54 Vangone A, Spinelli R, Scarano V, Cavallo L & Oliva R (2011) COCOMAPS: a web application to analyze and visualize contacts at the interface of biomolecular complexes. Bioinformatics 27, 2915–2916.

55 Oostenbrink C, Villa A, Mark AE & van Gunsteren WF(2004) A biomolecular force field based on the free enthalpy of hydration and solvation: the GROMOS force-field parameter sets 53A5 and 53A6. Journal of Computational Chemistry 25, 1656–1676.

56 Berendsen HJC, Postman JPM, van Gunsteren WF & Hermans J (1981) Interaction models for water in relation to protein hydration. In: Pullman B, editor. Intermolecular forces Dordrecht: Reidel Publishing Co. 331–342.

57 O’Boyle NM, Banck M, James CA, Morley C, Vandermeersch T & Hutchison GR (2011) Open Babel: An open chemical toolbox. Journal of Cheminformatics 3:33.

58 Trott O & Olson AJ (2010) AutoDock Vina: improving the speed and accuracy of docking with a new scoring function, efficient optimization and multithreading. Journal of Computational Chemistry 31, 455–461.

59 Morris GM, Huey R, Lindstrom W, Sanner MF, Belew RK, Goodsell DS & Olson AJ (2009) Autodock4 and AutoDockTools4: automated docking with selective receptor flexibility. Journal of Computational Chemistry 16, 2785–2791.

